# CRISPRi Screen Identifies a Novel Growth Suppressor lncRNA, *INSTAR*, in Human Monocytes

**DOI:** 10.1101/2025.10.30.685589

**Authors:** Christy Montano, Eric Malekos, Sergio Covarrubias, Sol Katzman, Lisa Sudek, Jillian Ward, Susan Carpenter

**Affiliations:** Department of Molecular, Cell and Developmental Biology, University of California Santa Cruz, California, USA; Department of Biomolecular Engineering, University of California Santa Cruz, California, USA

## Abstract

Acute myeloid leukemia (AML) is a heterogeneous disease that arises from dysregulated myeloid proliferation. We performed a high-throughput CRISPR interference (CRISPRi) screen in the THP-1 monocytic cancer cell line to identify long noncoding RNAs (lncRNAs) that play a role in contributing to cell proliferation. Our screen identified *INSTAR* (Intergenic Nuclear Suppressor lncRNA Targeting Adjacent Regulator *SFMBT2*) as a top candidate. RNA-seq on *INSTAR* deficient THP-1 cells revealed transcriptional changes in genes involved in cell proliferation as well as other cellular processes. Loss of *INSTAR* selectively reduced expression of its neighboring gene, *SFMBT2*. Functional assays confirmed that both genes suppress cell growth, revealing a cis-regulatory mechanism in which *INSTAR* regulates *SFMBT2* expression to control monocyte proliferation. Here, we leverage high-throughput screening to rapidly pinpoint functional lncRNAs providing novel insights into a key regulatory locus consisting of *INSTAR* and *SFMBT2* which could be critical for better understanding dysregulation contributing to acute myeloid leukemia.

## Introduction

According to the Leukemia and Lymphoma Society, leukemia is the most common cancer among children and adolescents.^1^ It has been reported that 1 in 4 children diagnosed with leukemia have acute myeloid leukemia (AML), which occurs when myeloid cells such as monocytes over-proliferate. AML is a heterogeneous disease with half of patients having large chromosomal abnormalities and the other half being cytogenetically normal.^2^ Currently, most newly diagnosed patients receive a chemotherapy regimen consisting of a combination of cytarabine and anthracycline which target DNA synthesis and disrupt the cell cycle.^3^ However, some patients might not be ideal candidates for this kind of therapy due to their unique genetic alterations, which lead to diverse molecular subtypes of the disease. These varying subtypes result in different treatment responses, which underscores the need for sensitive diagnostics and a tailored treatment plan for AML based on disease pathogenesis. Thus, research via the use of next-generation sequencing (NGS) has focused on identifying novel pathogenic targets that can accurately and non-invasively distinguish between these disease subtypes at initial diagnosis and recurrence.

RNA-sequencing has unveiled the surprising complexity of the human transcriptome. According to Gencode V49, over 35,000 long noncoding RNAs (lncRNAs) are expressed, which accounts for 46% of all annotated RNA.^4^ However, the vast majority do not yet have ascribed functions as molecular characterization approaches have lagged behind the speed of discovery.^4^ LncRNAs are transcripts that are more than 500 nucleotides in length and undergo both splicing and polyadenylation but lack coding potential.^5^ Beyond their cell-specific expression, recent research has highlighted their critical roles in immune cell signaling,^6-8^ differentiation,^9, 10^ as well as disease states including cancer. ^11-14^ A prime example from the cancer field is *PVT1* (Plasmacytoma Variant Translocation 1), one of the most well-known lncRNAs involved in tumor growth and metastasis.^11^ Studies have shown that the *PVT1* promoter has a tumor suppressor function *in cis*, by regulating the transcription of its neighbor, the crucial proto-oncogene *MYC*, a transcription factor that is essential for cell growth and division.^11^ In AML, the over-expression of the lncRNA *HOTTIP* (HOXA Distal Transcript Antisense RNA) increases expression of HOXA, which in turn results in altered hematopoietic stem cell self-renewal and differentiation, leading to leukemogenesis.^12^ According to Luo et al., *HOTTIP* may operate in a context-specific manner, acting either *in cis* or having trans functions outside of its locus particularly when it is overexpressed such as in AML.^12^ In cancers such as prostate cancer and B-cell lymphomas, lncRNAs including *GAS5* (Growth Arrest Specific 5), *HOTAIR* (HOX Transcript Antisense RNA), and *XIST* (X Inactive Specific Transcript) are differentially expressed, suggesting their potential as diagnostic markers as well as pathogenic targets.^13, 15^ Low expression of *GAS5* in plasma correlates with poor patient prognosis.^13^ In contrast, high *HOTAIR* expression is associated with cancer remission.^13^ Despite evidence that lncRNAs can influence monocyte proliferation, the full repertoire of lncRNAs that regulate monocyte proliferation and their underlying molecular mechanisms are still being defined.^5, 16-18^ To address this gap, we utilized the power of a high-throughput screen to identify functional candidate lncRNAs regulating monocyte proliferation.

We conducted a pooled CRISPR interference (CRISPRi) screen in human monocytic cells (THP-1).^19^ Utilizing a THP-1 specific lncRNA library targeting over 2,000 lncRNAs we identified a total of 16 growth suppressor gene hits. Following our preliminary RNA-seq analyses, we focused on our top hit *Linc02642* hereafter referred to as *INSTAR* (Intergenic Nuclear Suppressor lncRNA Targeting Adjacent Regulator *SFMBT2*). *INSTAR* was chosen for mechanistic studies due to three key factors: its ranking as our number one hit, its classification as an intergenic lncRNA with a distinct promoter, and its position adjacent to a protein-coding gene, *SFMBT2*, with a previously established role in cancer.^20, 21^ Knowing that lncRNAs can function *in cis,*^6, 11, 19, 22^ we sought to further explore *INSTAR*’s role as a cis regulator of *SFMBT2.* We found that *INSTAR* and *SFMBT2* lie within the same topologically associated domain (TAD) and that both of these transcripts reside in the chromatin compartment of THP-1 cells. We conducted functional experiments that validated *INSTAR* as a growth suppressor and revealed that loss of *SFMBT2* produced a similar phenotype. Altogether, this work demonstrates the importance of the *INSTAR* locus in regulating THP-1 cell proliferation, and underscores the value of high-throughput CRISPR screening for discovering functional lncRNAs.

## Materials and methods

### Cell lines

Wildtype (WT) THP1 cells were obtained from ATCC. All THP1 cell lines were cultured in RPMI 1640 supplemented with 10% low-endotoxin fetal bovine serum (ThermoFisher), 1X penicillin/streptomycin, and incubated at 37°C in 5% CO2. Cells were also treated with 100nM of PMA for 24 hr.

### Lentivirus production

All constructs were cotransfected into HEK293T cells with lentiviral packaging vectors psPAX (Addgene cat#12260) and pMD2.g (Addgene cat#12259) using Lipofectamine 3000 (ThermoFisher cat# L3000001) according to the manufacturer’s protocol. Viral supernatant was harvested 72h post-transfection as previously described.^36^

### THP1-NFkB-EGFP-dCasKRAB

We constructed a GFP-based NF-κB reporter system by adding 5x NF-κB-binding motifs (GGGAATTTCC) upstream of the minimal CMV promoter-driven EGFP. THP1s were lentivirally infected and clonally selected for optimal reporter activity. Reporter cells were then lentivirally infected with the dCas9 construct that was constructed using Lenti-dCas9-KRAB-blast, Addgene #89567. Cells were clonally selected for knockdown efficiency greater than 90%.

### THP1-NfKB-EGFP-dCASKRAB-sgRNA

NFkB-EGFP-CRISPRi-THP1 cells were lentivirally infected with sgRNAs. sgRNA constructs were made from a pSico lentiviral backbone driven by an EF1a promoter expressing T2A flanked genes: puromycin resistance and mCherry as previously described.^45^ sgRNAs were expressed from a mouse U6 promoter. Twenty-nucleotide forward/reverse gRNA oligonucleotides were annealed and cloned via the AarI site.

### SCREENING PROTOCOL

#### sgRNA library design and cloning

10 sgRNAs were designed for each TSS of hg19 annotated lncRNAs expressed in THP1s at baseline and upon stimulation. The sgRNA library also included 700 non-targeting control sgRNAs, and sgRNAs targeting 50 protein-coding genes as positive controls. The sgRNA library was designed and cloned as previously described.^6^

#### Screening and analysis

THP1-CRISPRi cells were infected with the sgRNA genome-scale library at a low multiplicity of infection (MOI = 0.3). Three days post-infection, cells were puromycin-selected (10 μg/ml) for 5 days to obtain cherry-positive (sgRNA) cells and were maintained at >1000X coverage at all times. Cells were grown for a total of 21 days and maintained at over 1000X coverage throughout. Genomic DNA was extracted from triplicate day 21 samples, and libraries prepared and sequenced as previously described.^6^ For the growth screen, the Day 21 sample was compared to the plasmid library pool. fastq.gz files were analyzed using the gRNA_tool: https://github.com/quasiben/gRNA_Tool. All guide RNA (sgRNA) were collapsed to obtain raw sgRNA counts. Counts were normalized to the median, and fold-changes were calculated for each sgRNA. To identify significant genes for the growth screen, the Mann-Whitney U test was performed comparing fold-changes for sgRNAs targeting each gene to non-targeting controls previously described.^24^ Data is available at GSE305555.

#### Sequencing Data

RNA sequencing was performed to compare negative control THP-1 cells to those with *INSTAR*, *STARD7-AS1,* or *SNHG17* knockdown. Data are available at GSE305555.

#### RNA Isolation and RT-qPCR

Cells were homogenized in Tri-Reagent (Sigma Aldrich, T9424-200 mL). RNA was extracted with the Direct-zol RNA miniprep plus RNA extraction Kit (Zymo, R2072). 1 ug of total RNA was reverse transcribed into cDNA (iScript cDNA synthesis kit, Bio-Rad cat# 1708840). cDNA was diluted 1:3 in qPCR experiments. QPCR (iTaq SYBRgreen Supermix, Bio-Rad cat# 1725121) was run using the cycling conditions as follows: 50C for 2 min, 95 °C for 2 min followed by 40 cycles of 95 °C for 15s, 60 °C for 30s, and 72 °C for 45s.

#### TOPO-TA Cloning

Isoforms were cloned using the Life Technologies TOPO-TA kit protocol and reagents (Thermo cat# K457501). Different isoform sequences are available in Table 1.

**Table 1.**
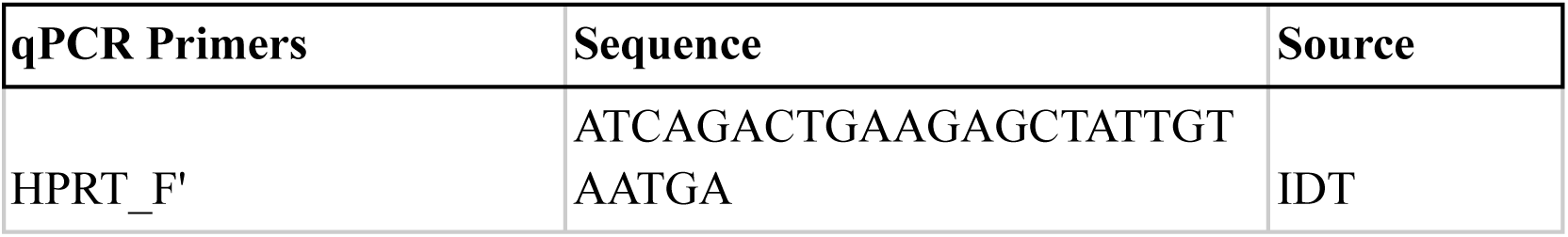
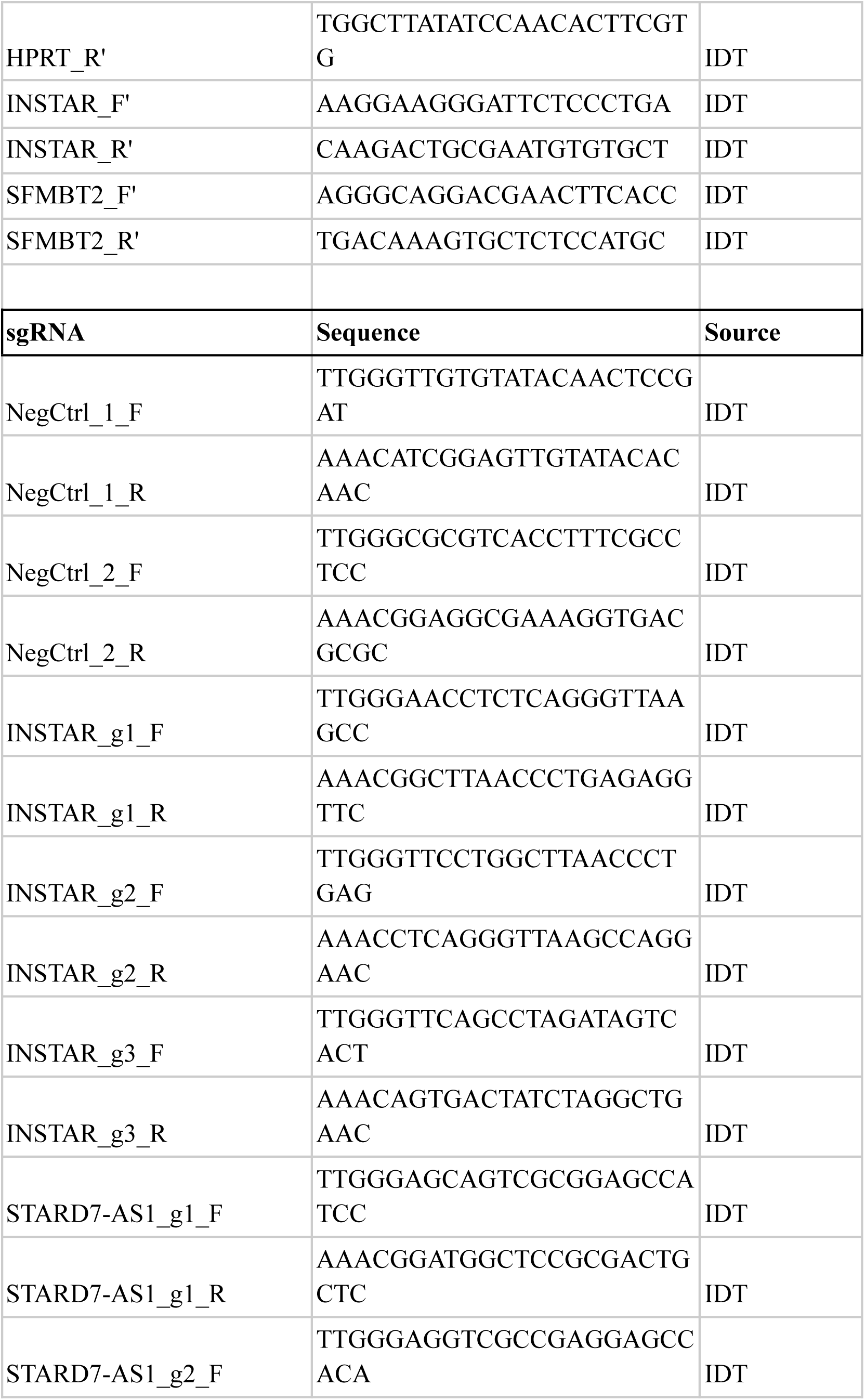
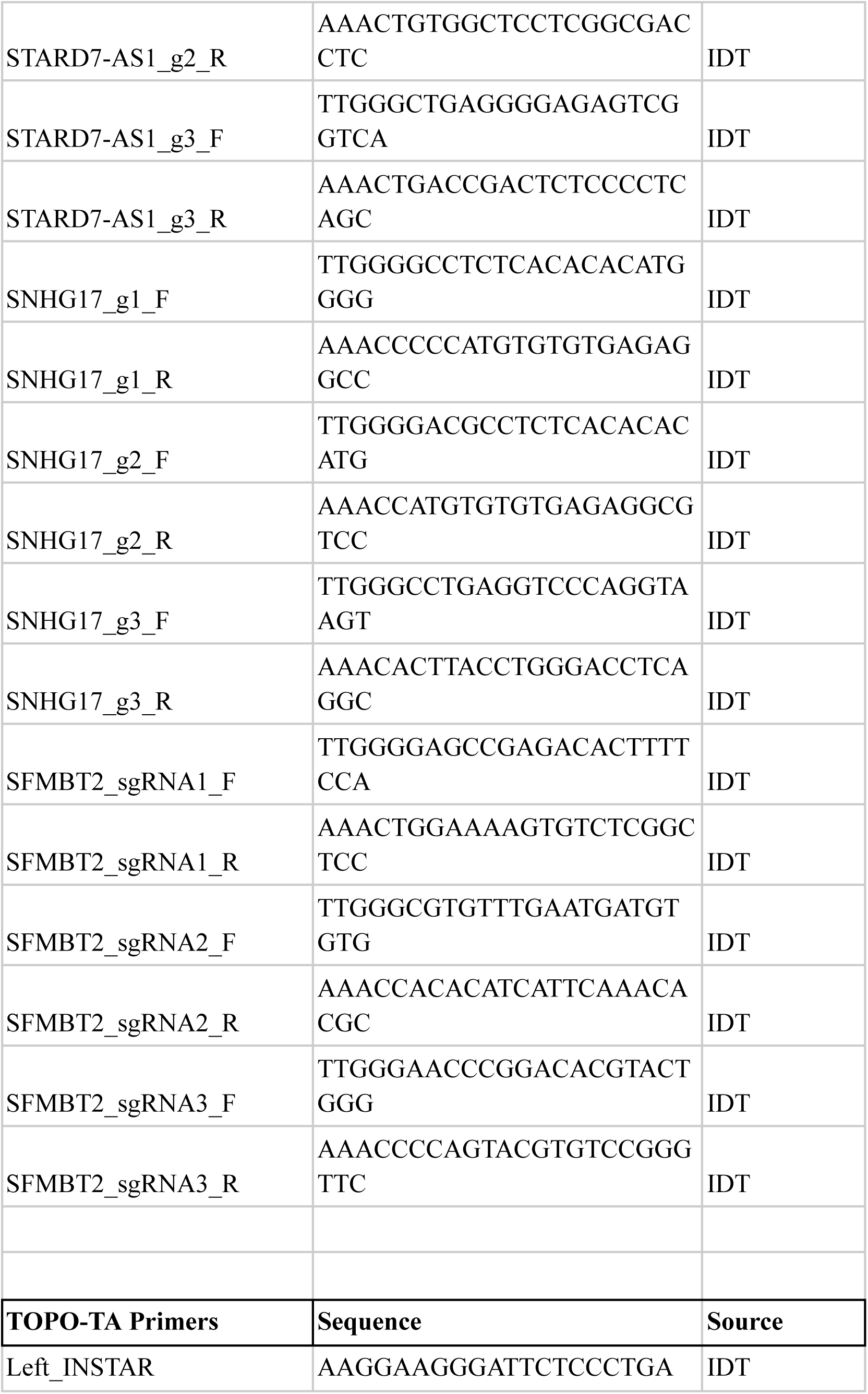
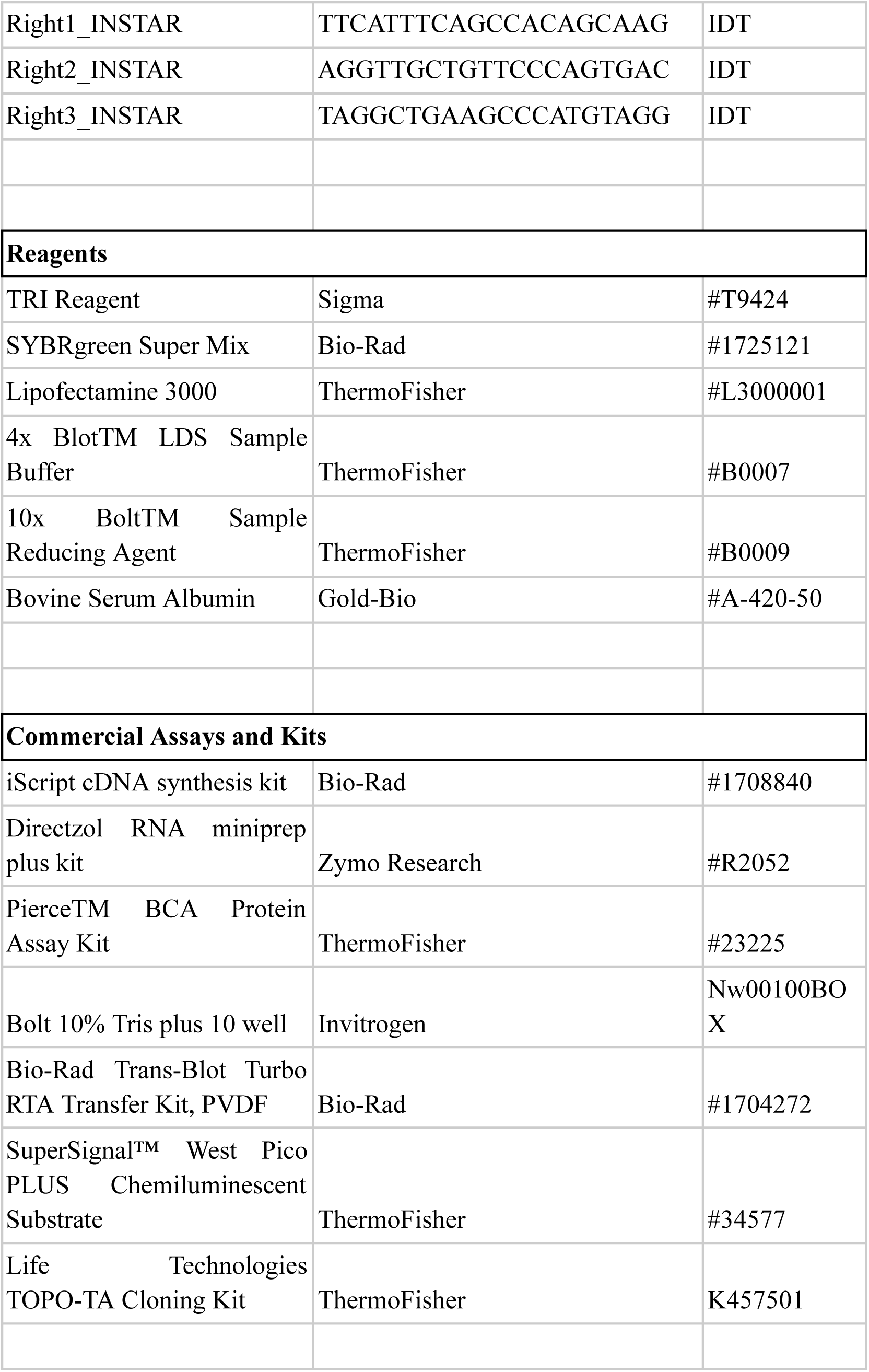
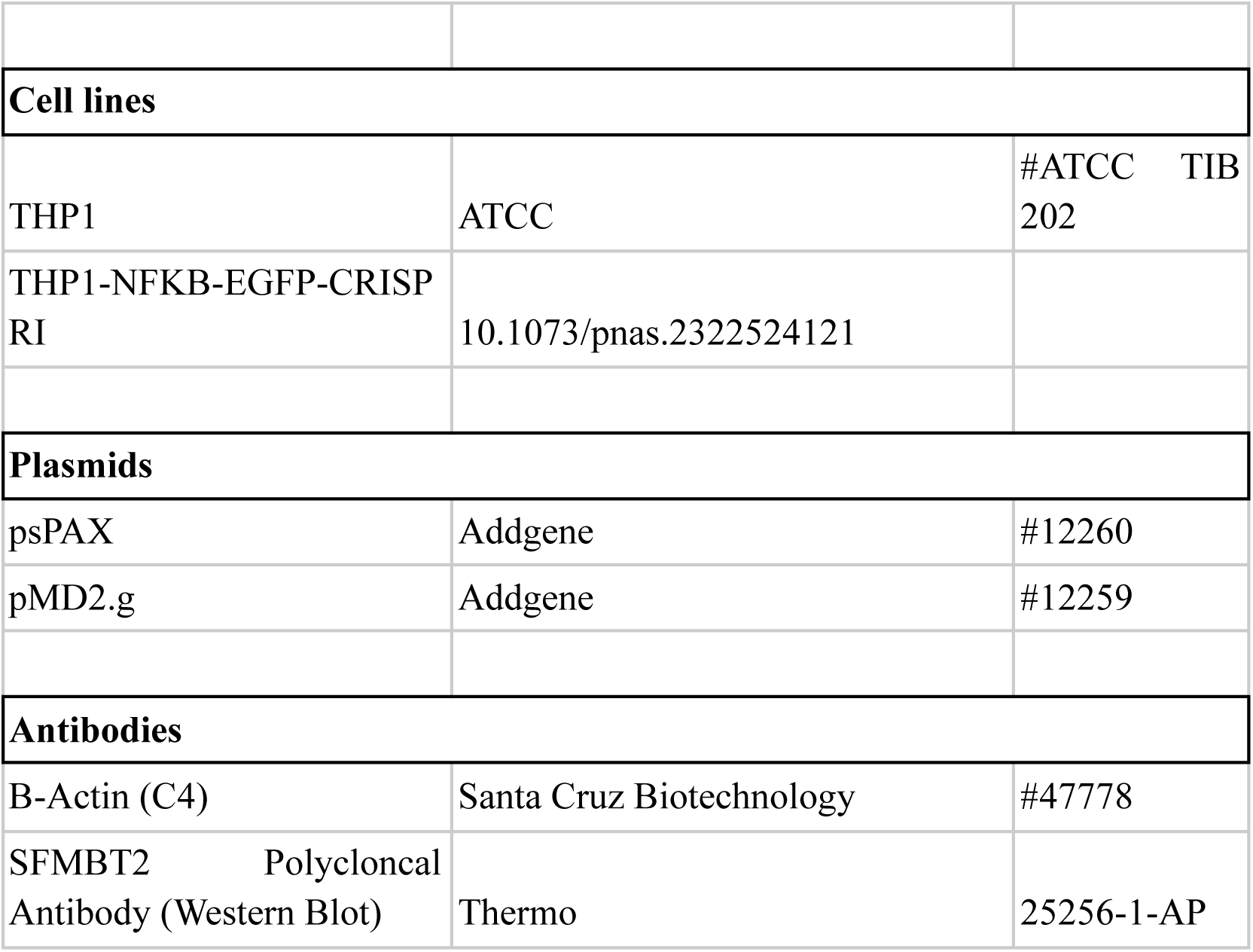
Primers and Reagents.

#### Mixed cell proliferation assay

We used the top three performing sgRNA guides from the screen to validate the lncRNA genes. Briefly, sgRNA-infected THP1 cells (cherry-pos) were mixed with uninfected THP1 cells (cherry-neg) at a 1:1 ratio in triplicate. We used flow cytometry to monitor the ratio of cherry-pos to cherry-neg cells at 0- and 35-days post-plating. All validation cytometry was performed on the Attune NxT Flow Cytometer.

#### Nuc/Cyt fractionation and RT-qPCR

WT THP1 cells were fractionated according to the NE-PER kit protocol (ThermoFisher cat# 78833) with RNAse inhibitor (Superase-IN, ThermoFisher cat# AM2696) added to the cytosolic and nuclear lysis buffers. 3 volumes of Trizol (TRI Reagent, Sigma T9424) were added to the fractions, and RNA was isolated using the Direct-zol RNA Miniprep Plus Kit (Zymo cat# R2052). 16uL of RNA isolated from fractions was reverse transcribed (iScript cDNA synthesis kit, Bio-Rad cat# 1708840), followed by qPCR (iTaq SYBRgreen Supermix, Bio-Rad cat# 1725121) using the cycling conditions as follows: 50C for 2 min, 95 °C for 2 min followed by 40 cycles of 95 °C for 15s, 60 °C for 30s, and 72 °C for 45s.

#### Cellular Fractionation and Isolation of Chromatin-Associated RNA

Followed the protocol as described by Conrad and Ørom.^46^ 16uL of RNA isolated from fractions was reverse transcribed (iScript cDNA synthesis kit, Bio-Rad cat# 1708840), followed by qPCR (iTaq SYBRgreen Supermix, Bio-Rad cat# 1725121) using the cycling conditions as follows: 50C for 2 min, 95 °C for 2 min followed by 40 cycles of 95 °C for 15s, 60 °C for 30s, and 72 °C for 45s.

#### Western Blot

Cell lysates were prepared by removing cell media, washing cells with 1× PBS, and then lysing them in RIPA buffer (150 mM NaCl, 1.0% Nonidet P-40, 0.5% sodium deoxycholate, 0.1% SDS, 50 mM Tris-HCL [pH 7.4]. And 1.0 mM EDTA) containing protease inhibitor cocktail (Roche, cat#04693124001). Protein in lysates was quantified using the PierceTM BCA Protein Assay Kit (Thermo Fisher cat#PI23227). Equal amounts of protein (38ug) of each sample were denatured at 70 °C for 10 min prior to loading on 10% SDS-PAGE. Samples were transferred to polyvinylidene difluoride (PVDF) membranes using the Standard protocol on the Trans-Blot Turbo Transfer System (Bio-Rad). Membranes were blocked with TBST (1× Tris-buffered saline with 0.1% Tween 20), supplemented with 5% (wt/vol) nonfat dry milk for 1 h and blotted with primary antibody SFMBT2 (1:1000, Proteintech Catalog Number: 25256-1-AP) at 4 °C overnight. Horseradish peroxidase (HRP)-conjugated goat anti-rabbit (1:2,000, Bio-Rad, #1706515) secondary antibody was used andWestern blots were developed with SuperSignalTM West Pico PLUS Chemiluminescent Substrate (Thermo Scientific cat# 34577). Horseradish peroxidase-conjugated β-actin (1:500, Santa Cruz Biotechnology, sc-47778) was used as loading control. Prior to running controls, deactivation of HRP by 0.2% sodium azide was performed for 2 h. Control blots were developed using Pierce ECL (Life Technologies, 32106). ImageJ (rsbweb.nih.gov/ij/) analysis was performed to quantify band intensities for each Western blot.

## Results

### CRISPRi Screen Identifies Growth Suppressor lncRNAs in Human Monocyte THP-1 Cells

We previously conducted a dropout CRISPRi screen to identify lncRNAs that act as viability and growth suppressors in THP-1 cells.^19^ While our viability hits were characterized previously,^19^ here we focus on functional characterization of those lncRNAs involved in proliferation. We infected THP-1 cells with a pooled lentivirus library. This library was designed with 10 sgRNAs per lncRNA, specifically targeting the transcription start site (TSS) of over 2,000 annotated lncRNAs (GRCh37, hg19) (Fig. 1A). Following infection, we selected THP-1 cells with puromycin for 7 days. To identify sgRNAs that either dropped out or enriched over time, we extracted genomic DNA after 21 days (Fig. 1A). To rank hits we compared the fold-change of TSS-targeting sgRNAs to non-targeting control sgRNAs, assigning each a Mann-Whitney U-score (Fig. 1B, C). Genes with U-scores greater than 3 were classified as growth suppressors (Fig. 1C). Positive controls included *PVT1*, *CASP8AP2*, and *OLMALINC*, previously characterized as viability or growth suppressor lncRNAs.^11, 19, 23^ In total, 16 growth suppressor lncRNAs met this threshold with p-adjusted values of less than 0.1 (Fig. 1B, C).

**Figure 1:**
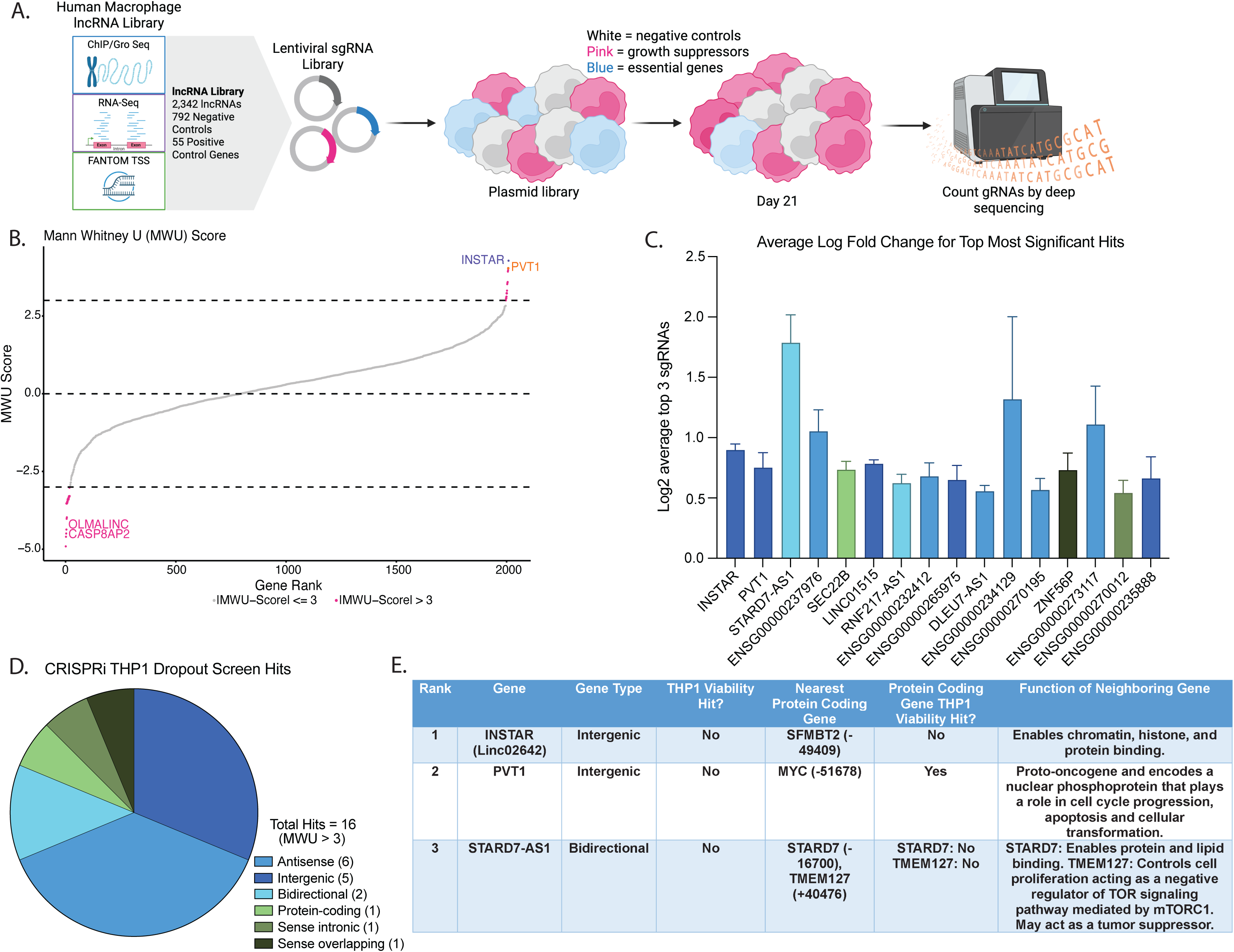
CRISPRi Screen identifies growth-suppressor lncRNAs in human THP-1 monocytic leukemia cells. **(A) Overview of CRISPRi THP-1 dropout screen.** LncRNA transcription start sites (TSSs) were predicted using ChIP-Gro-Seq, RNA-sequencing, and FANTOM TSS data. NFkB-EGFP-CRISPRi-THP1 cells were subsequently infected with a pooled sgRNA library comprising over 2,000 sgRNAs targeting Gencode hg19 annotated lncRNA TSSs. After puromycin selection, samples were collected on day 21, and sgRNAs from the original plasmid library and day 21 were PCR amplified and sequenced. (B) CRISPRi THP-1 dropout screen analysis. Mann-Whitney U analysis was performed on each of the three screen replicates to determine sgRNA enrichment. Genes with MWU scores of -3 and 3 were considered significant. (C) Significant growth suppressor hits. The average log2 fold-change of the top three best-scoring sgRNAs for all significant hits with SD. Hits are color-coded to match their gene category in D. (D) Dropout screen hit categories. Significant hits were categorized based on gene/ lncRNA type using the UCSC Genome Browser (Human hg38). (E) Top 3 hits table. This table summarizes the top 3 lncRNA growth suppressor hits, *INSTAR* and *STARD7-AS1*, and the positive control hit, *PVT1*.

Given the nature of CRISPRi machinery and its ability to bidirectionally silence transcription within ∼1kb of the target sgRNA,^24^ it is unsurprising that some of the top hits identified were antisense or bidirectional lncRNAs (Fig. 1C, D), a trend we also observed in other screens. ^5, 6, 10^ Heterochromatin spreading likely extended to these protein-coding genes, leading to their knockdown alongside the lncRNAs. For example, *STAG2-AS1* (ENSG00000232412) neighbors *STAG2*, a protein-coding gene with established involvement in tumorigenesis, cell proliferation, and growth.^25^ Similarly, *STARD7-AS1* (ENSG00000204685), a bidirectional hit, shares a promoter with *STARD7* (Fig. 1C), which regulates placental cancer cell proliferation and migration.^26^ The second most prominent lncRNA hit category identified were intergenic lncRNAs, with promoters at least 1kb away from their neighboring protein-coding genes (Fig. 1D, E). One of our hits, *SEC22B*, represents a protein-coding gene, previously known to be involved in plasma cell proliferation and maintenance.^27^ Lastly, 2 of our hits were found to be overlapping their nearest protein-coding gene. Interestingly, none of their neighboring protein-coding genes have been previously identified as regulators of cell proliferation; potentially making them novel coding regulators of this process (Supp. Table 1).

To determine which candidate to pursue further, we performed preliminary RNA-seq analyses on 3 lncRNAs, initially chosen based on their rank and proximity to proliferation-associated genes. These include *STARD7-AS1, SNHG17,* and a previously uncharacterized lncRNA we named, Intergenic Nuclear Suppressor lncRNA Targeting Adjacent Regulator *SFMBT2* (*INSTAR).* We cloned the top 3 performing guides of *STARD7-AS1* and our preliminary RNA-seq results showed that we achieved successful knockdown of *STARD7-AS1*; however, due to its proximity to its protein-coding gene neighbor we also observed significant knockdown of *STARD7* (Fig. S1). Gene Ontology (GO) analysis of the differentially expressed genes (DEG) showed that the most significant genes (|L2FC| > 1.5) were involved in cell development, differentiation, and proliferation (Fig. S1). We also cloned the top 3 performing guides of *SNHG17*, however because these guides targeted the most common isoforms and not their specific transcription start sites (TSS) we only achieved about 50% knockdown (Fig. S2). As a result, GO analysis of DEGs could only be performed on the most significantly upregulated genes (L2FC > 1.5) which revealed that they were involved in cell metabolic processes (Fig. S2). In addition, we cloned the top 3 performing guides of our top hit, *INSTAR* (Fig. 2A). GO analysis on the most significant DEGs showed their involvement in cell proliferation, development, and metabolism (Fig. 3B, E). Since *INSTAR* was our top intergenic hit, successfully knocked down, and associated with proliferation-related GO terms, we prioritized it for further mechanistic investigation.

**Figure 2:**
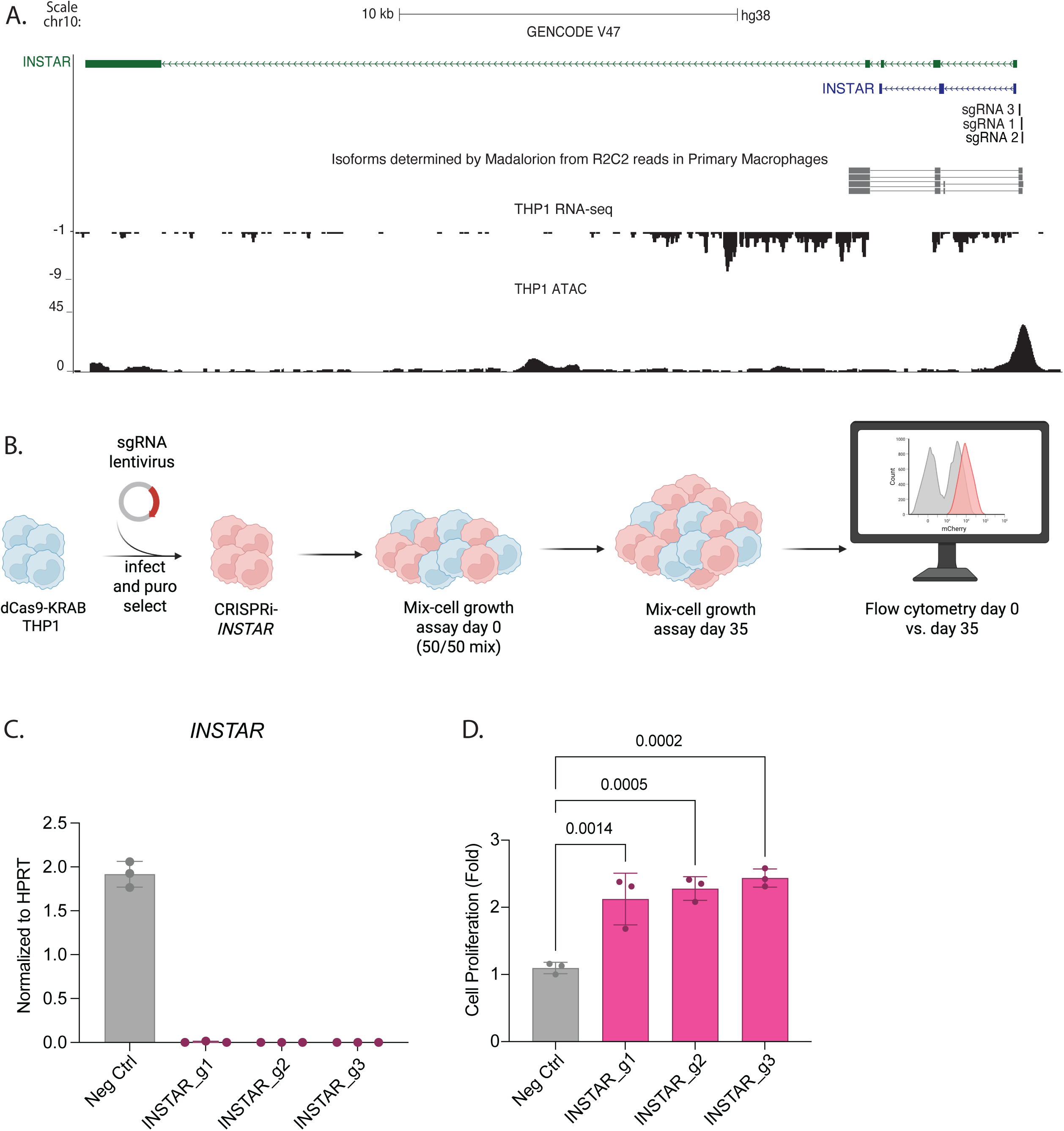
Functional validation of *INSTAR* as a growth suppressor in THP-1 cells. **(A) sgRNAs targeting *INSTAR*.** The browser track displays the TOPO-TA determined dominant isoform of *INSTAR* in THP-1 cells in blue and the 3 sgRNAs (sgRNA 1, sgRNA 2, and sgRNA 3) designed to knock down *INSTAR*. Browser tracks include: isoforms in primary human macrophages,^28^ RNA-sequencing (GSE150571), and ATAC-sequencing reads (GSE96800) of the *INSTAR* locus in wildtype THP-1 cells. **(B) Overview of THP-1 cell proliferation assay.** The top three sgRNAs were cloned into a plasmid with an mCherry fluorescent reporter, followed by lentivirus infection and puromycin selection of dCas9-KRAB THP1 cells. Mcherry-positive cells were mixed with mcherry-negative cells in a 1:1 ratio and plated in triplicate for each sgRNA for each gene. Cells were flowed every week for a total of 35 days. **(C) CRISPRi knockdown of *INSTAR* in THP-1 cells.** RT-qPCR of *INSTAR* across three biological replicates shows a statistically significant knockdown of *INSTAR* by all three sgRNAs vs. a non-targeting sgRNA (Neg Ctrl). Values are normalized to HPRT. **(D) THP-1 cell competition assay results for *INSTAR*-KD.** We monitored the proliferation of sgRNA-edited (mCherry-positive) cells compared to unedited (mCherry-negative) cells over 35 days, starting with a 1:1 mixture. The experiment was repeated 3 times, and a representative experiment is shown. Rate of change was calculated as the percentage difference in the averaged fold change between the knockdown cell lines and the non-targeting control.

**Figure 3:**
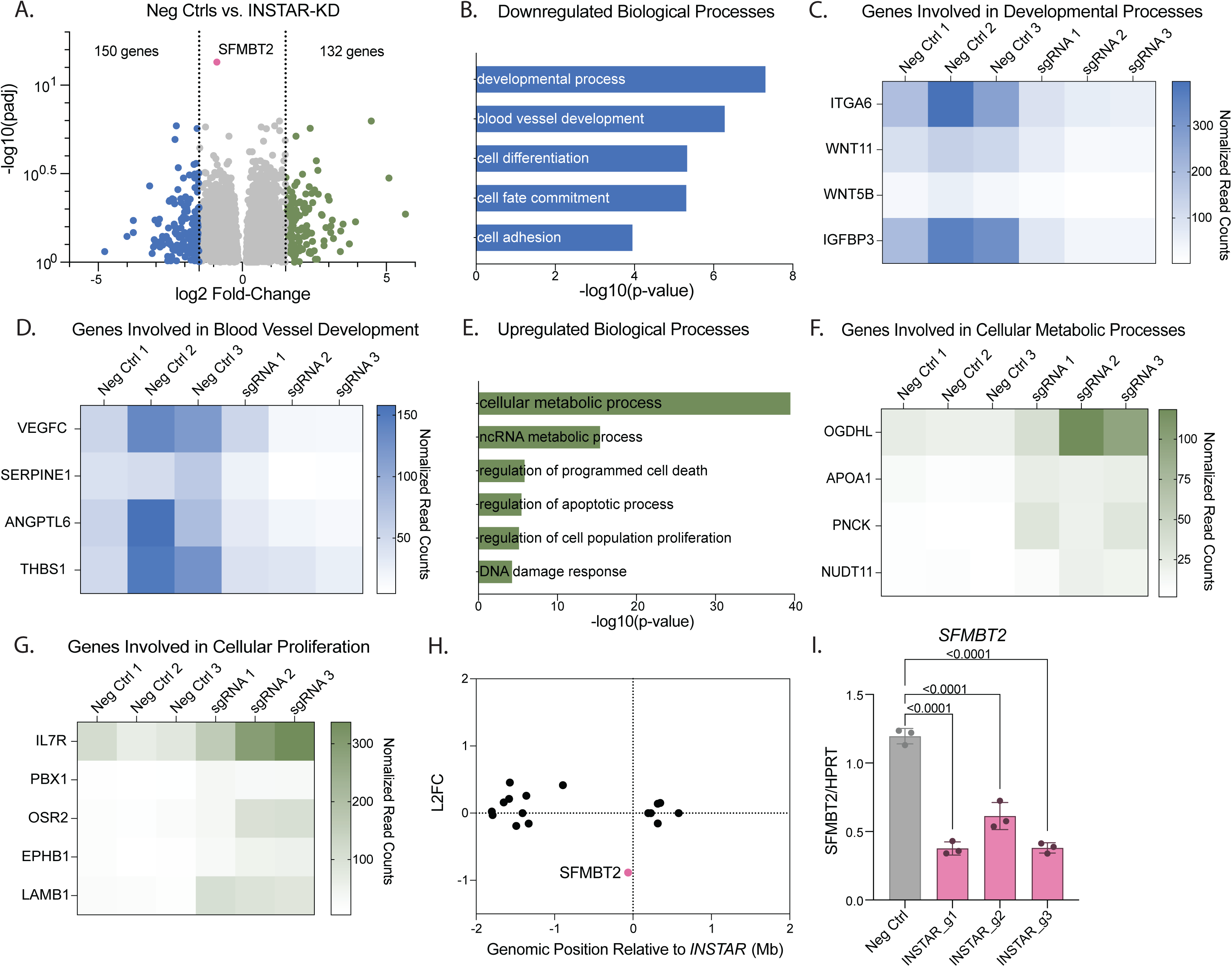
RNA-Seq reveals reduced *SFMBT2* expression following *INSTAR* knockdown. **(A) Negative controls vs. *INSTAR-*KD RNA sequencing analysis.** DESeq2 was used to establish the log2 fold-change of genes between three independent negative control sgRNAs and three sgRNAs targeting *INSTAR* to identify upregulated and downregulated genes upon CRISPRi *INSTAR* knockdown. |L2FC| > 1.5 were considered significant. (B) Enrichment analysis of downregulated genes in *INSTAR*-KD THP-1 cells. Enriched GO terms of downregulated genes (L2FC < -1.5) after knockdown of *INSTAR.* (C-D) Heat maps of downregulated developmental genes in *INSTAR* knockdown cells. Heat maps show normalized read counts of significantly downregulated genes in *INSTAR* knockdown cells (n=3) compared to negative controls (n=3). Genes are grouped by their involvement in (C) cell developmental processes and (D) blood vessel development. (E) Enrichment analysis of upregulated genes in *INSTAR*-KD THP-1 cells. Enriched GO terms of upregulated genes (L2FC > 1.5) after knockdown of *INSTAR*. (F-G) Heat maps of upregulated metabolic and proliferative genes in *INSTAR*-KD knockdown cells. Heat maps show normalized read counts of significantly upregulated genes in *INSTAR* knockdown cells (n=3) compared to negative controls (n=3). Genes are grouped by their involvement in (F) cell metabolic processes and (G) cell proliferation. (H) *INSTAR*-KD reveals *SFMBT2* as the most statistically significant downregulated gene. Differential gene expression of genes in the local region (± 2 Mb) of *INSTAR*, depicted as L2FC between three independent negative control sgRNAs and three sgRNAs targeting *INSTAR*. (I) Expression of *SFMBT2* in *INSTAR*-KD cells. Expression of *SFMBT2* was measured in *INSTAR* deficient cells by RT-qPCR in each of 3 CRISPRi cell lines (sgRNA 1, sgRNA 2, sgRNA 3) along with a nontargeting control (Neg Ctrl). Values are normalized to HPRT and error bars represent standard deviation. This graph represents one of three independent experiments with similar results.

### Functional Validation of *INSTAR* as a Growth Suppressor in THP-1 Cells

*INSTAR* (*ENSG00000232591*), emerged as the top hit from our CRISPRi THP-1 dropout screen with an adjusted p-value of less than 0.1 and a MWU-score of 4.3. This intergenic lncRNA is situated on chromosome 10, approximately 50kb beyond the promoter of the protein-coding gene, *SFMBT2* (Scm-like with Four MBT Domains 2). ATAC-seq revealed epigenetic signatures indicative of chromatin accessibility at its TSS, an important feature for designing effective sgRNAs, as CRISPR targeting efficiency is enhanced in open chromatin regions (Fig. 2A). Although *INSTAR* is annotated with five isoforms averaging 28kb (GRCh38/hg38), according to the Genotype-Tissue Expression (GTEx) Project, *INSTAR* is a lowly expressed lncRNA, with a median expression of less than 1TPM across tissues (Fig. S3). Consistent with this, our WT THP-1 RNA-seq data also showed *INSTAR* to be lowly expressed and did not provide sufficient resolution to determine the dominant isoform (Fig. 2A). Given that lncRNAs are poorly annotated, it was important to determine the exact isoform of *INSTAR* in our cells of interest, THP-1s. Our analysis using isoform-specific primers (Table 1), TOPO-TA cloning followed by PCR identified a predominant 4.5kb isoform with three exons in THP-1 cells (Fig. 2A, blue isoform). These results were corroborated by our primary human macrophage isoform long-read nanopore data, further highlighting lncRNA isoform specific cell-type expression (Fig. 2A).^28^

To further test *INSTAR*’s potential role as a growth suppressor lncRNA, we chose the top three performing sgRNAs from our CRISPRi screen according to their adjusted p-values and cloned them into an mCherry reporter plasmid to generate individual lines lacking the lncRNA (Fig. 2A, B). Using RT-qPCR we confirmed knock down of the *INSTAR* transcript by more than 90% (Fig. 2C). To validate our screen result and confirm *INSTAR* had a growth suppressor phenotype we conducted a mixed-cell proliferation assay (Fig. 2B). For this assay we combined mCherry positive cells (sgRNA containing) with mCherry negative cells at a 1:1 ratio and monitored cell proliferation over time as measured by flow cytometry (Fig. 2B). All three sgRNAs targeting *INSTAR* resulted in more than 80% increased cell proliferation compared to the negative control sgRNA (Fig. 2D). This cell proliferation assay confirmed that *INSTAR* plays a role in THP-1 cell proliferation.

### RNA-Seq Reveals Reduced *SFMBT2* Expression Following *INSTAR* Knockdown

To identify genes and pathways impacted by *INSTAR* knockdown during cell proliferation, we performed RNA-seq and subsequent gene enrichment analyses. We used three non-targeting controls and three sgRNAs targeting *INSTAR*. This approach identified 282 differentially expressed genes (DEGs) in *INSTAR*-deficient cells, with an absolute log2-fold change (L2FC) of ≥1.5 (Fig. 3A). Gene Ontology (GO) analysis of these DEGs revealed key biological pathways influenced by *INSTAR*. The most significantly downregulated genes (L2FC < -1.5) were enriched in developmental processes (e.g., I*TGA6, WNT11, WNT5B, IGFBP3*), blood vessel development (e.g., *VEGFC, SERPINE1, ANGPTL6, THBS1*), cell differentiation, cell fate commitment, and cell adhesion (Fig. 3B, C, D). Conversely, the most significantly upregulated genes (L2FC > 1.5) were associated with cellular metabolic processes (e.g., *OGDHL, APOA1, PNCK, NUDT11*), noncoding RNA metabolic processes, regulation of cell death, regulation of apoptotic process, regulation of cell proliferation (e.g., *IL7R, PBX1, OSR2, EPHB1, LAMB1*), and DNA damage response (Fig. 3E, F, G).

Interestingly, the most statistically significant hit from our RNA-seq data was *SFMBT2*. It was downregulated with an L2FC of -0.9 and a p-adj < 0.001 (Fig. 3A). *SFMBT2* was the only DEG with a significant adjusted p-value and L2FC within a 2-Mb window of *INSTAR* (Fig. 3H). We confirmed this finding using RT-qPCR, showing a ∼50% decrease in *SFMBT2* expression in CRISPRi-*INSTAR* cells (Fig. 3I). Collectively, these results indicate that *INSTAR* is an intergenic growth suppressor lncRNA that regulates key biological processes, including proliferation, development, and metabolism.

### *INSTAR*-Mediated Regulation of Monocyte Proliferation through *SFMBT2*

We next examined the mechanism by which *INSTAR* regulates *SFMBT2* transcript expression by first analyzing the chromatin and epigenetic landscape of the *INSTAR* locus. Using previously published high-throughput chromosome conformation capture (Hi-C) data from THP-1 cells,^29^ we found that *INSTAR* and *SFMBT2* occupy the same topologically associating domain (TAD), a finding supported by CTCF ChIP-seq and quantified chromosome interaction frequencies (Fig. 4A). The presence of the active chromatin mark H3K27ac on the *INSTAR* promoter further suggests its potential for transcription factors to engage and trigger transcription at this site (Fig. 4A). To determine where *INSTAR* and *SFMBT2* RNA localize within THP-1 cells, we performed cellular fractionation and RT-qPCR, which showed that both *INSTAR* and *SFMBT2* transcripts are predominantly nuclear (>90%) (Fig. 4B). Furthermore, isolation of chromatin-associated RNA revealed that both transcripts are largely bound to chromatin (Fig. 4C).

**Figure 4:**
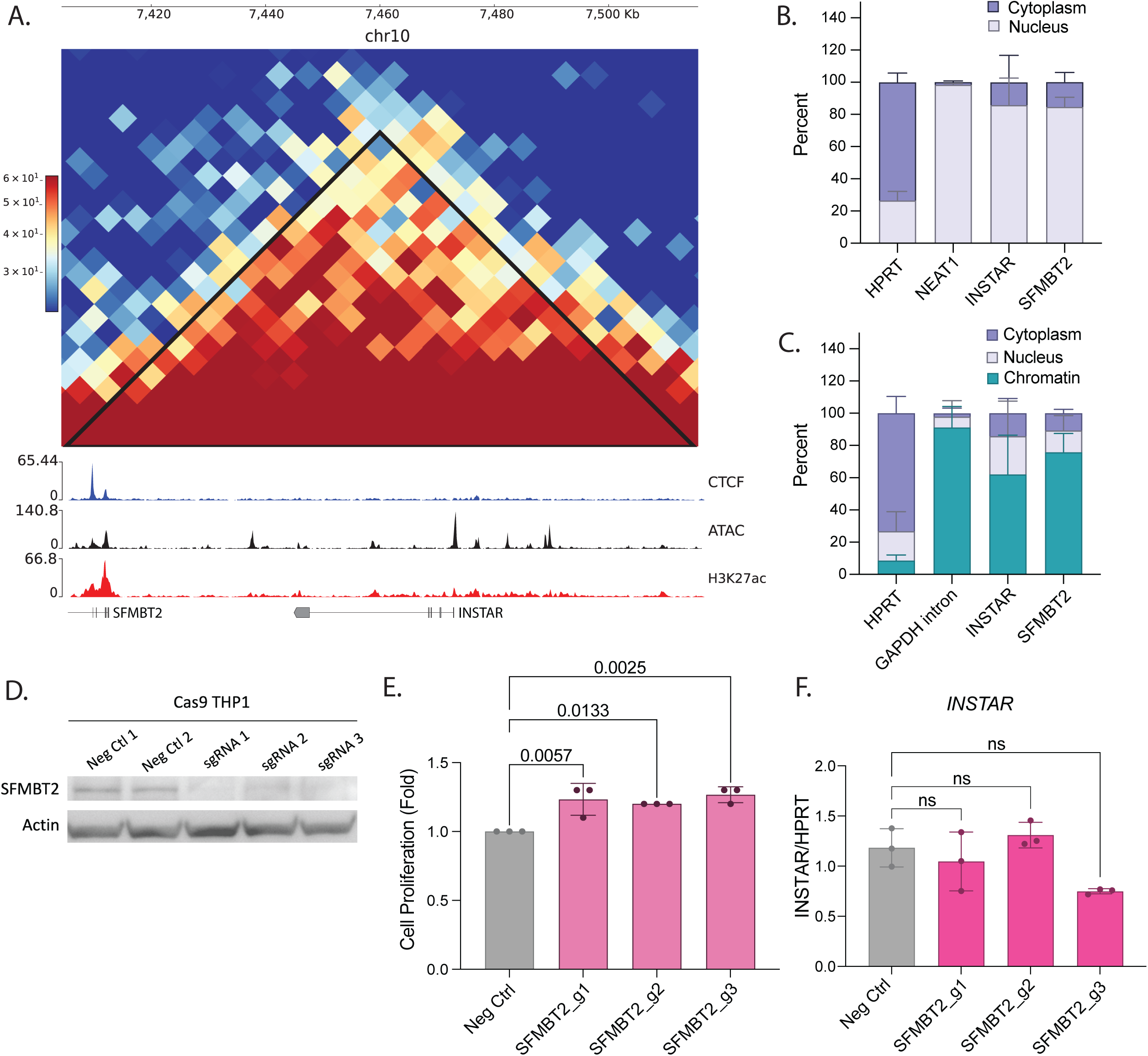
*INSTAR*-mediated regulation of monocyte proliferation through *SFMBT2*. **(A) Predicted TAD in the *INSTAR* locus in THP-1 cells.** Browser tracks display regions of CTCF binding, ATAC peaks, and regions of H3K27 acetylation. Numbers on the y-axis represent raw read counts. **(B-C) Cellular localization of *INSTAR* in THP-1 cells.** (B) Nuclear/ cytoplasmic fractionation of wildtype THP-1 cells. Error bars represent standard deviation across three biological experiments. (C) Chromatin fractionation of wildtype THP-1 cells. Error bars represent standard deviation across three biological experiments. **(D) Western blot analysis of *SFMBT2*.** Negative control vs. 3 *SFMBT2* knockdown cell lines (sgRNAs 1-3). **(E) THP-1 cell competition assay results for *SFMBT2*-KD cells.** We monitored the proliferation of sgRNA-edited (mCherry-positive) cells compared to unedited (mCherry-negative) cells over 35 days, starting with a 1:1 mixture. Error bars represent standard deviation across three biological experiments. **(F) Expression of *INSTAR* in *SFMBT2*-KD cells.** Expression of *INSTAR* was measured in *SFMBT2* deficient cells by RT-qPCR in each of 3 CRISPR-Cas9 cell lines (sgRNA 1, sgRNA 2, sgRNA 3) along with a nontargeting control (Neg Ctrl). Values are normalized to *HPRT* and error bars represent standard deviation. This graph represents one of three independent experiments with similar results.

Given that *INSTAR* knockdown significantly reduced *SFMBT2* expression (Fig. 2I) and both genes reside within the same TAD (Fig. 4A),^29^ a domain where DNA regulatory elements influence gene expression, we tested whether the growth suppressor phenotype could also be driven by *SFMBT2*. Instead of CRISPRi we employed CRISPR-Cas9 with the goal of causing a frame-shift and knockout of the *SFMBT2* protein using three distinct sgRNAs cloned into an mCherry reporter vector, and knockdown was confirmed by Western blot (Fig. 4D). Our mixed-cell proliferation assays demonstrated that *SFMBT2* deficiency increases THP-1 cell proliferation (20-30%) (Fig. 4E). Notably, RT-qPCR of the Cas9-*SFMBT2* cell lines showed no significant change in *INSTAR* expression compared to our negative control sgRNA (Fig. 4F). Together these findings suggest that *INSTAR* impacts THP-1 cell proliferation by regulating its protein-coding gene neighbor, *SFMBT2*, *in cis*.

## Discussion

The heterogeneity of AML poses a major challenge for diagnosis and treatment, highlighting the need to define its underlying molecular mechanisms.^30, 31^ Recent efforts have focused on lncRNAs, a large class of transcripts whose cell-type specific expression makes them ideal candidates for elucidating disease mechanisms. Given the recognized role of monocytes and macrophages in AML pathogenesis,^32^ investigating the lncRNAs that regulate these immune cells provides a promising avenue for identifying novel biomarkers and therapeutic targets.^18,17^ Evidence supports a role for lncRNAs in monocyte proliferation; however, the breadth of lncRNAs involved and the molecular pathways through which they function remain unclear.^5, 16-18^ To address this gap, we have utilized the power of CRISPR screening to identify functional lncRNAs that regulate THP-1 cell proliferation, focusing on the top hit, *INSTAR*. Notably, *INSTAR* was included in Lui et al.’s screening library but was not identified as a hit in their large-scale screening studies, further underscoring lncRNA cell type specificity.^5^ Remarkably, even within the subset of lncRNAs expressed in seven cell types, over 80% of hits were unique to one.^5^ By identifying *INSTAR* as a growth suppressor lncRNA, our findings emphasize the importance of cell-specific approaches for characterizing lncRNA function.

Having validated that *INSTAR* knockdown increases THP-1 cell proliferation (Fig. 2C-D), we next asked whether it can function *in cis* to regulate neighboring genes or move away from its native locus to impact genes *in trans*. Following RNA-seq there were many genes with altered expression upon *INSTAR* knockdown but we were intrigued to find that *SFMBT2,* the only gene within a 2-Mb window of the *INSTAR* locus, showed the most significant downregulation (Fig. 3H-I). *INSTAR* and *SFMBT2* reside within the same TAD in THP-1 cells (Fig. 4A), prompting us to test whether *SFMBT2* mediates the observed growth suppressor phenotype. We found that knocking out *SFMBT2* led to a 20% increase in cell proliferation compared to our negative control cell line, demonstrating that *SFMBT2* also has a growth suppressor phenotype in THP-1 cells (Fig. 4E). *SFMBT2* exhibits cell-specific roles across cancers. While the protein has been reported to act as a growth suppressor,^20, 21, 33^ the circular isoform circ-*SFMBT2* shows oncogenic activity.^34^ These contrasting effects likely reflect differences in protein regulation and circ-*SFMBT2* formation or function. Additionally, *SFMBT2* has emerged as a proliferative hit in screens from JHOC-5, hTERT-RPE1, and K562 cells.^35^ However, *SFMBT2* was not a hit on previous screens in THP-1 cells or in our own mouse macrophage screen.^35, 36^ Polycomb group proteins (PcG) such as *SFMBT2* remodel chromatin to coordinate developmental gene expression.^37^ *SFMBT2* acts as a transcriptional repressor, but its function appears context dependent, potentially driving tumor-suppressive or oncogenic programs depending on the cellular context and target genes. ^21, 33, 34^ This type of context-specific behavior has also been reported for other PcG members.^38^

In mice *SFMBT2* is essential for proper trophoblast development, and its loss leads to embryonic lethality.^39, 40^ Phenotypic data from the IMPC showed that female mice heterozygous for *SFMBT2* showed elevated monocyte counts, indicating that *SFMBT2* dosage is important for myeloid homeostasis (Fig. S4).^40^ In line with this we showed that *INSTAR* knockdown increased THP-1 proliferation by ∼80% (Fig. 2D) despite only a 50% decrease of *SFMBT2* mRNA (Fig. 3I). Indeed we only observed a weak protein signal for *SFMBT2* by Western blot (Fig. 4D) and it is dominantly retained as an mRNA in the nucleus bound to chromatin (Fig. 4C). *SFMBT2*’s function appears critically dependent on its protein levels. We propose that THP-1 cells manage this dosage post-transcriptionally through nuclear retention of *SFMBT2* mRNA. This mechanism would act as a buffer, holding transcripts at the chromatin to control their release for translation and dampen gene expression noise from transcriptional bursts, which is a strategy previously described by Bahar Halpern et al.^41^ Given that *SFMBT2* is a powerful repressor, tight regulation of its own expression is crucial for sustaining polycomb-mediated repression at target loci. This is especially relevant in monocytes and macrophages, which require tight regulation of immune genes. For example, in prostate cancer, the loss of *SFMBT2* protein is linked to the aberrant, NFkB-driven activation of inflammatory chemokines.^20, 21^ Therefore, it is logical that immune cells like THP-1s would employ multiple control mechanisms, such as nuclear retention, to keep this strong repressor in check and ensure a precise inflammatory response.

Further work will be needed to better understand the specific expression levels of *SFMBT2* and its association with AML and other human diseases. While analysis of publicly available datasets reveals no significant correlation between *SFMBT2* expression and overall survival in AML (Fig. S5), its role in leukemogenesis warrants further investigation. Notably, *SFMBT2* is highly expressed in granulocyte-macrophage progenitors (GMPs) during healthy (normal) hematopoiesis (Fig. S6) and exhibits elevated and variable expression across AML subtypes (Fig. S7).^42^ Future molecular studies are therefore essential to elucidate the precise mechanistic contributions of both *SFMBT2* and *INSTAR* to the pathogenesis of AML. It is unclear whether *INSTAR* is functionally conserved in mice, as our analyses revealed that *INSTAR* lacks both syntenic and sequence conservation. While the protein itself, *SFMBT2*, is conserved between humans and rodents there are a number of distinct regulatory differences. The main one is rodents harboring a cluster of miRNAs within intron 10, shown to be critical for the regulation of the protein.^43^

Although *INSTAR* is not conserved at a sequence level, there is evidence that *SFMBT2* can be regulated *in cis* by a lncRNA as Engreitz et al. reported a cis-regulatory mechanism between *Sfmbt2* and *Blustr* (*linc1319*) in mouse embryonic stem cells.^44^ This is interesting in the context of critical protein coding genes between species undergoing tight regulation by neighboring lncRNAs. A recent study reported that haploinsufficiency of the lncRNA *CHASERR* (CHD2 Adjacent Suppressive Regulatory RNA) increases *CHD2*, a dosage-sensitive transcription factor whose dysregulation leads to neurodevelopmental disorders.^22^ Although *CHD2* and *SFMBT2* have distinct functions, notably, both are essential nuclear proteins whose expression is tightly controlled through cis-regulatory interactions with neighboring lncRNAs. These parallels suggest that other lncRNAs may similarly regulate nearby protein-coding genes *in cis*, and that disrupting this feedback could contribute to disease.

Our study shows that the *INSTAR* locus plays a significant role in controlling THP-1 cell proliferation by regulating its neighbor *SFMBT2 in cis*. Although we learned that *INSTAR* and *SFMBT2* act as growth suppressor genes and localize to the chromatin compartment it is yet to be determined whether the lncRNA transcript or the DNA regulatory elements present at the locus are responsible for the observed growth suppressor phenotype. Previous research shows that for some examples the lncRNA transcript is not necessary for them to locally regulate their protein-coding neighbor *in cis*.^11, 44^ In these cases, it has been shown that the processes associated with lncRNA production, such as transcription and splicing, enable gene regulation by bringing promoter and enhancer regulatory elements into close proximity.^11, 44^ In other cases only a single copy of the lncRNA is required for the gene to function as an enhancer RNA and to regulate its neighbor *in cis.*^7^ Future experiments could focus on exploring this mechanism by using Cas13 technology, which binds and cleaves RNA. Whereas CRISPRi silences gene loci through heterochromatin formation, sometimes unintentionally affecting nearby protein-coding genes or regulatory elements, Cas13 would enable specific perturbation of the *INSTAR* transcript to assess its effects on *SFMBT2* expression and THP-1 cell phenotype. A recent study utilized Cas13 to screen for lncRNAs functioning as viability and growth regulators, unfortunately *INSTAR* was not targeted in their library.^16^ CRISPRi and Cas13 together provide powerful tools to decipher the many complex ways in which lncRNAs can function to regulate biological processes. Together with recent studies, our results demonstrate the importance of cis-acting lncRNA mechanisms in the regulation of complex signaling pathways and the power of CRISPR screening to identify functional genes for mechanistic validation.

## Acknowledgements

We thank Dr. Alice Devigne of the UCSC CRISPR Core for their services. We thank Dr. Ravindra Majeti for his helpful discussions on AML and assistance in utilizing publicly available resources on hematopoiesis.

## Author Contributions

C.M performed the validation and downstream molecular mechanistic experiments of hits from the screens and RNA-seq approaches. S.Co performed the CRISPR based screen. E.M, S.K performed the bioinformatic analysis to support the RNA-seq and screening studies. L.S performed and analyzed Western blot in Fig. 4. J.W helped with analyzing RT-qPCR experiments in Fig. 3. C.M and S.C conceived and coordinated this project. C.M and S.C wrote the manuscript with input from all coauthors.

## Data availability

All data utilized in this study is available through the GEO portal on NCBI.

- RNA-seq was performed on THP1-NFkB-EGFP-CRISPRi-sgRNA controls (nontargeting and anti-GFP) versus CRISPRi-lncRNA knockdown in monocytes (THP1s). Data are available at GSE305555.
- Data pertaining to ATACSeq and ChIPSeq in THP1s are available at GEO: GSE96800 and SRA: PRJNA385337.
- Isoforms from primary human macrophage processed data can also be explored as a UCSC Genome Browser session at https://genome.ucsc.edu/s/vollmers/IAMA.

