## Supplementary Table and Figure Legends for "CRISPRi Screen Identifies a Novel Growth Suppressor lncRNA, *INSTAR*, in Human Monocytes"

**Supplementary Table 1: Extended Significant Hits Table.** Supplemental table with information on all significant (MWU < 3) hits. Gene information was gathered using the UCSC Genome Browser (GRCh38/ hg38), The Human Gene Database, and BioGRID Open Repository of CRISPR Screens (ORCS).

**Figure S1: Knockdown of *STARD7-AS1* leads to changes in cell development and proliferation processes. (A) sgRNAs targeting *STARD7-AS1.*** The browser track displays the 3 sgRNAs (sgRNA 1, sgRNA 2, and sgRNA 3) designed to knock down *STARD7-AS1*. Browser tracks include: isoforms in primary human macrophages,[28] RNA-sequencing (GSE150571), and ATAC-sequencing reads (GSE96800) of the *STARD7-AS1* locus in wildtype THP-1 cells. **(B) Negative controls vs. *STARD7-AS1*-KD RNA sequencing analysis.** DESeq2 was used to establish the log2 fold-change of genes between three independent negative control sgRNAs and three sgRNAs targeting *STARD7-AS1* to identify upregulated and downregulated genes upon CRISPRi *STARD7-AS1* knockdown. |L2FC| > 1.5 were considered significant. **(C-D) Normalized Read Counts from DESeq Analysis.** Normalized read counts from DESeq analysis for *STARD7-AS1* and (D) its neighboring protein-coding gene *STARD7*. **(E) Enrichment analysis of downregulated genes in *STARD7-AS1*-KD THP-1 cells.** Enriched GO terms of upregulated genes (L2FC > 1.5) after knockdown of *STARD7-AS1*. **(D) Enrichment analysis of upregulated genes in *STARD7-AS1*-KD THP-1 cells.** Enriched GO terms of upregulated genes (L2FC > 1.5) after knockdown of *STARD7-AS1*.

**Figure S2: Knockdown of *SNHG17* leads to changes in cell development and proliferation processes. (A) sgRNAs targeting *SNHG17.*** The browser track displays the 3 sgRNAs (sgRNA 1, sgRNA 2, and sgRNA 3) designed to knock down *SNHG17*. Browser tracks include: isoforms in primary human macrophages,[28] RNA-sequencing (GSE150571), and ATAC-sequencing reads (GSE96800) of the *SNHG17* locus in wildtype THP-1 cells. **(B) Negative controls vs. *SNHG17*-KD RNA sequencing analysis.** DESeq2 was used to establish the log2 fold-change of genes between three independent negative control sgRNAs and three sgRNAs targeting *SNHG17* to identify upregulated and downregulated genes upon CRISPRi *STARD7-AS1* knockdown. |L2FC| > 1.5 were considered significant. **(C) Normalized Read Counts from DESeq Analysis.** Normalized read counts from DESeq analysis for *SNHG17*. **(D) Enrichment analysis of upregulated genes in *SNHG17*-KD THP1 cells.** Enriched GO terms of upregulated genes (L2FC > 1.5) after knockdown of *SNHG17*.

**Figure S3: Genotype Tissue Expression (GTEx) of *INSTAR.*** Expression of *INSTAR* in different tissues according to GTEx (V10) RNA-seq of 19788 samples (946 donors) across 54 tissues. Numbers on the y-axis represent transcripts per million (TPM). The data used for this figure were obtained from: the GTEx Portal on 10/02/2025 and/or dbGaP accession number phs000424.vN.pN on 10/02/2025.

**Figure S4: Homeopoetic Phenotype of *Sfmbt2*$^{em1(IMPC)Mbp}$ mouse from the International Mouse Phenotyping Consortium (IMPC).** Each point represents statistical annotations for their given phenotype as represented by the label on the y-axis. The dotted line represents a manual annotation by the IMPC, at $1x10^{-15}$ in order to be displayed in the chart. According to the MMRRC mice were generated as follows: CRISPR guide(s) and Cas9 protein were microinjected or electroporated into C57BL/6NCrl zygotes and progeny were screened for the desired mutation. Founders were mated to C57BL/6NCrl breeders, and derived N1 progeny were identified by PCR and/or sequencing. N1 were then mated again to C57BL/6NCrl breeders, to generate N2 mice, identified by PCR and/or sequencing. N2 heterozygous mutant mice were mated for production of phenotyping cohorts, and N2 or N2F1 heterozygous mice were used for cryopreservation purposes. The data used for this figure were obtained from: www.mousephenotype.org on 10/03/2025.

**Figure S5: RNA-seq expression of SFMBT2 in Human Acute Myeloid Leukemia Samples from The Cancer Genome Atlas (TCGA).** Kaplan-Meier plot generated using tcga-survival.com. Kaplan-Meier curves are generated by dividing patients into two groups and comparing the survival times between each group. For RNA-Seq, microRNAs, and RPPA, the division is made based on the mean expression of the feature. For RNA-Seq the division is made based on the mean expression of SFMBT2.

**Figure S6: Hierarchical tree that shows SFMBT2 gene expression in different healthy hematopoietic cells.** Expression levels are visualized according to their color codes and the size of the node indicates the median of the cell type. Human HSC cells are from GSE17054. Human GMP, MEP cells are from GSE19599. Human Monocytes cells are from GSE11864. Human Monocytes cells are from E-MEXP-1242. Data has been batch corrected. The data used for this figure were obtained from: BloodSpot 3.0 a database of gene and protein expression data in normal and malignant haematopoiesis.[42]

**Figure S7: Hierarchical tree that shows SFMBT2 gene expression in different healthy vs AML hematopoietic cells.** Expression levels are visualized according to their color codes and the size of the node indicates the median of the cell type. Human Normal Hematopoiesis cells are from GSE42519. Human AML cells are from GSE13159. Data has been batch corrected. The data used for this figure were obtained from: BloodSpot 3.0 a database of gene and protein expression data in normal and malignant haematopoiesis.[42]
