## Supplementary Table and Figures for "CRISPRi Screen Identifies a Novel Growth Suppressor lncRNA, *INSTAR*, in Human Monocytes"

### Supplementary Table 1

| Rank | Gene | Function | Type | THP1 Cell Proliferation Hit?* | Nearest Neighbor | Promoter Distance (bp) | Neighbor Function | Neighbor THP1 Cell Proliferation Hit?* |
|---|---|---|---|---|---|---|---|---|
| 1 | ENSG00000232591 | Unknown | Intergenic | N/A | SFMBT2 | -49409 | Enables chromatin, histone, and protein binding. | No |
| 2 | ENSG00000249859 | Transcription of this gene is regulated by tumor suppressor p53. | Intergenic | N/A | MYC | -58239 | Proto-oncogene | No |
| 3 | ENSG00000204685 | Unknown | Bidirectional | N/A | STARD7 | -16700 | Enables protein and lipid binding. | No |
| 4 | ENSG00000237976 | Unknown | Antisense | N/A | RFX5 | -246 | Enables DNA-binding TF activity, specific to RNA-polymerase II. | No |
| 5 | SEC22B | The protein encoded by this gene is a member of the SEC22 family of vesicle trafficking proteins. | Protein-coding | N/A | N/A | N/A | N/A | N/A |
| 6 | ENSG00000228065 | Nasopharyngeal carcinoma and autism. | Intergenic | N/A | LRRT3 | >1 Mb | May play a role in the development and maintenance of the vertebrate nervous system. | No |

| | | | | | | | | |
|---|---|---|---|---|---|---|---|---|
| 7 | ENSG00000236548 | Unknown | Bidirectional | N/A | RNF217 | -47302 | E3 ubiquitin protein ligase, transfers ubiquitin to targeted substrates. | No |
| 8 | ENSG00000232412 | Unknown | Antisense | N/A | STAG2 | -232 | Required for the cohesion of sister chromatids after DNA replication. | Yes |
| 9 | ENSG00000265975 | Unknown | Intergenic | N/A | PIK3R5 | -6552 | Required for the recruitment of the catalytic subunit to the plasma membrane. | Yes |
| 10 | ENSG00000237152 | Associated with leukemia. | Antisense | N/A | RNASEH2B | -81223 | Degrades the RNA of RNA:DNA hybrids. | Yes |
| 11 | ENSG00000234129 | Associated with Rhabdoid Tumor Predisposition. | Antisense | N/A | HCCS | 53 | The protein encoded by this gene is an enzyme that covalently links a heme group to the apoprotein of cytochrome c. | No |
| 12 | ENSG00000270195 | Unknown | Antisense | N/A | SLBP | -493 | This gene encodes a protein that binds to the stem-loop structure in replication- | Yes |

| | | | | | | | | |
|---|---|---|---|---|---|---|---|---|
| | | | | | | | dependent histone mRNAs. | |
| 13 | ENSG00000267419 | Unknown | Pseudogene (sense-overlapping) | N/A | ZNF506 | 15987 | May be involved in transcriptional regulation. | Yes |
| 14 | ENSG00000273117 | Associated with autism. | Antisense | N/A | INSIG1 | -1047 | Enables protein, lipid, and oxysterol binding. | No |
| 15 | ENSG00000270012 | Unknown | Sense intronic | N/A | PPP1R3F | 4557 | Enables protein and glycogen binding. | No |
| 16 | ENSG00000235888 | Associated with colorectal cancer. | Intergenic | N/A | PSMG1 | 120000 | Enables molecular adaptor activity. Involved in chaperone-mediated protein complex assembly. | Yes |

*BioGRID Open Repository of CRISPR Screens (ORCS)

### Supplementary Figure (S1)

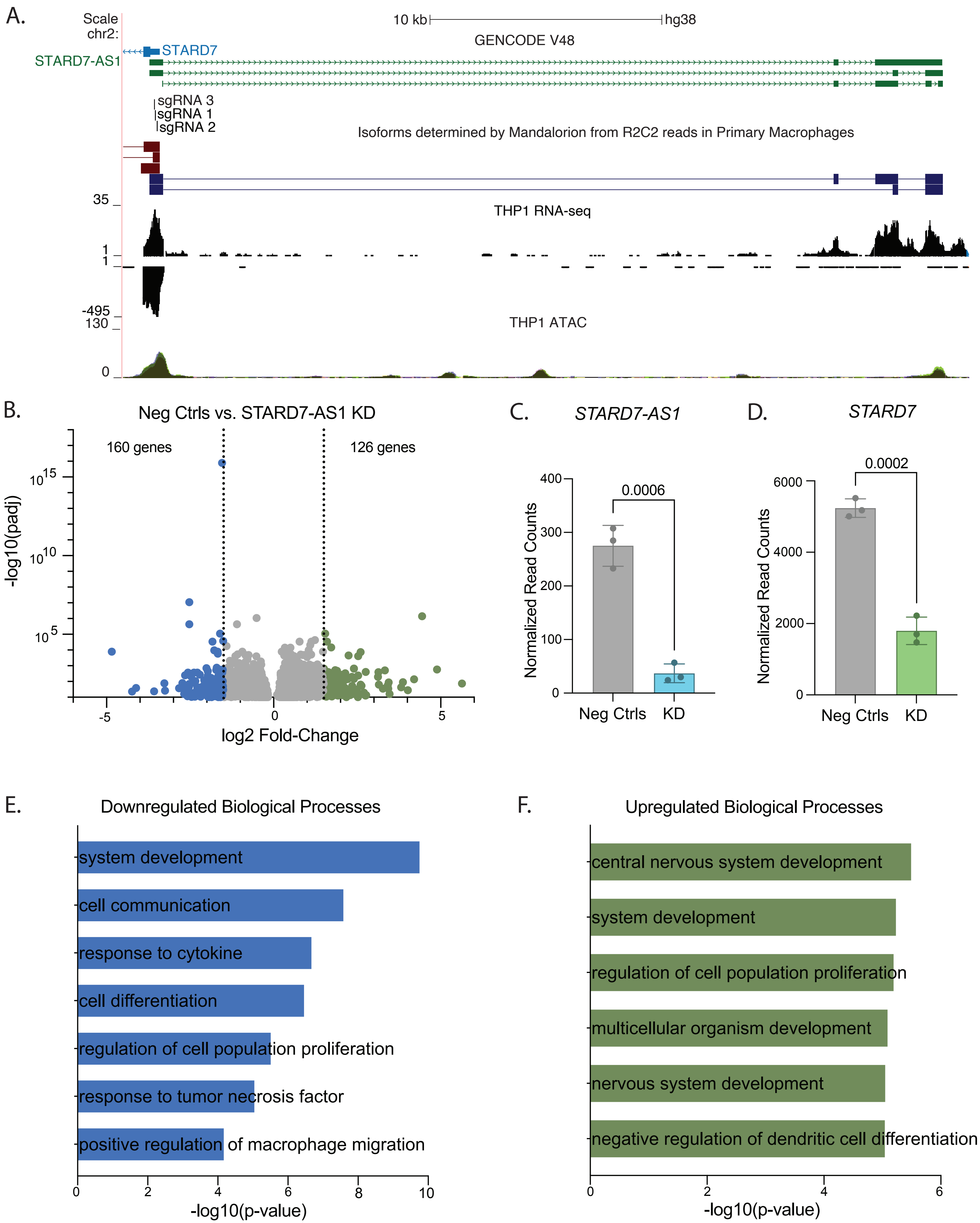

### Supplementary Figure (S2)

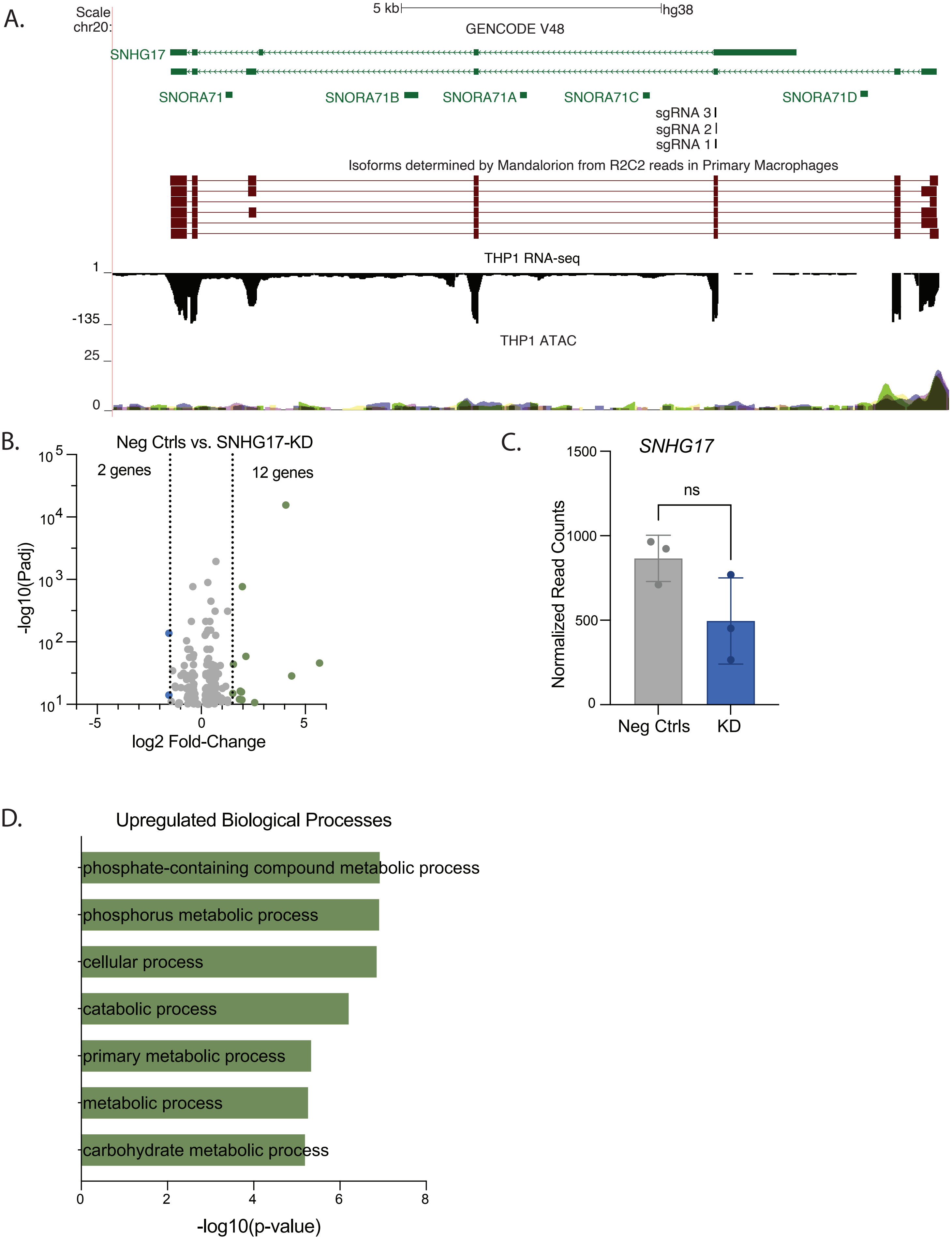

### Supplementary Figures (contd.)

S3.

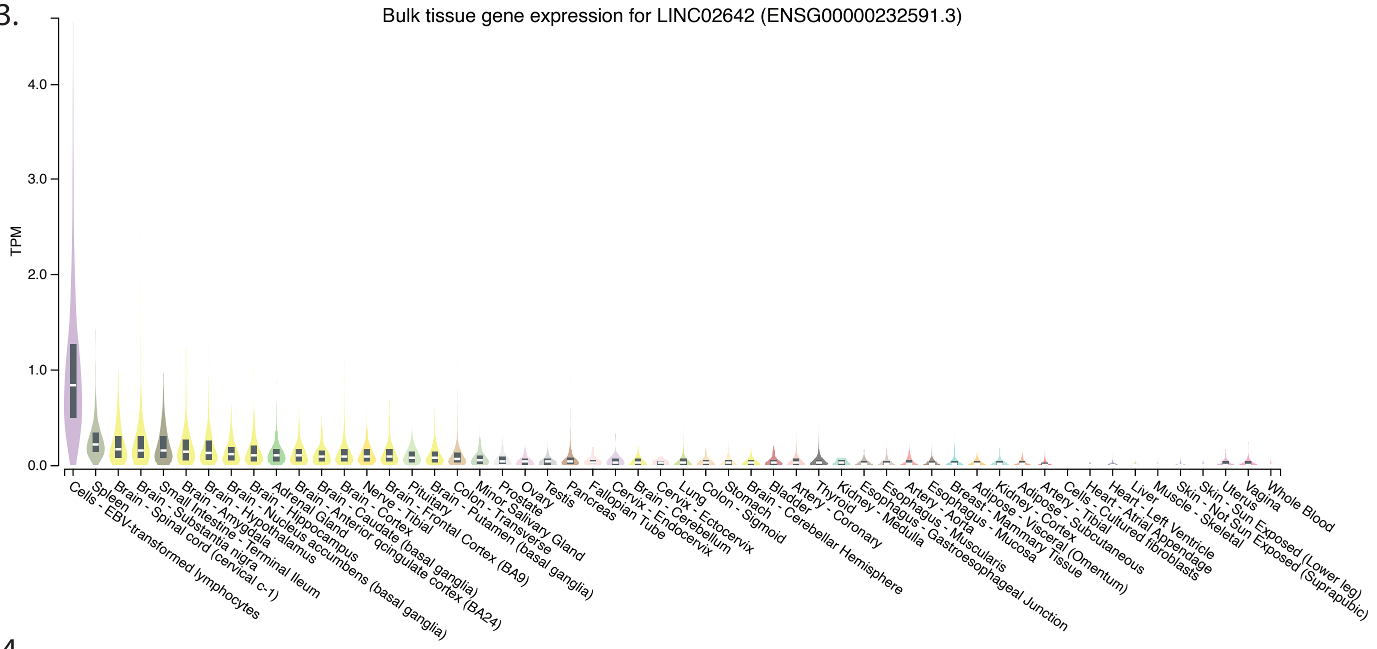


S4.

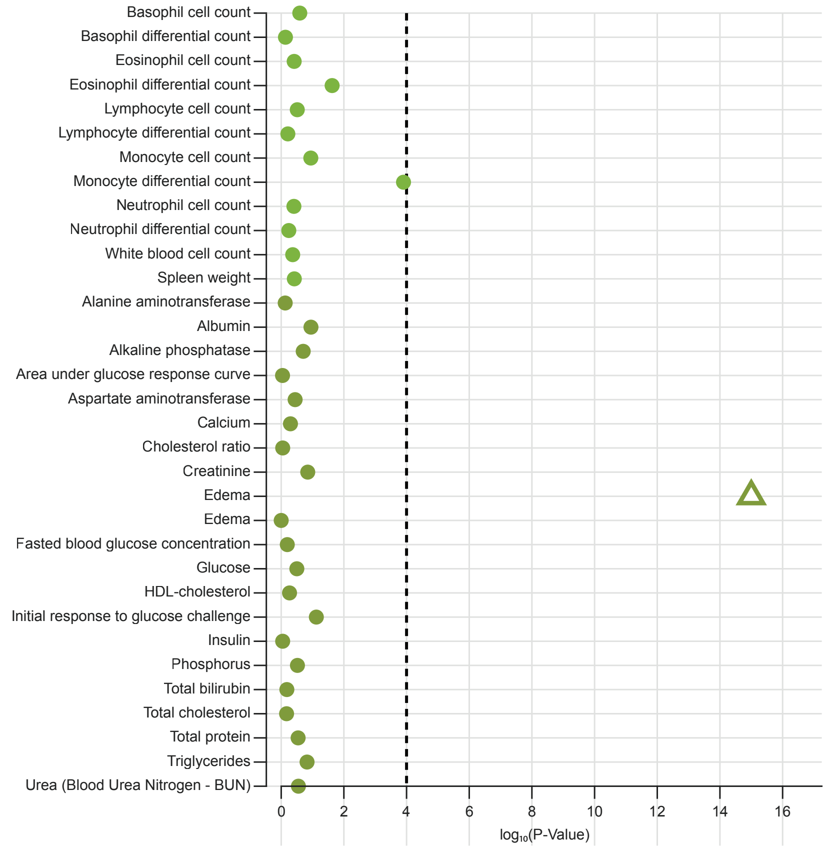


S5.

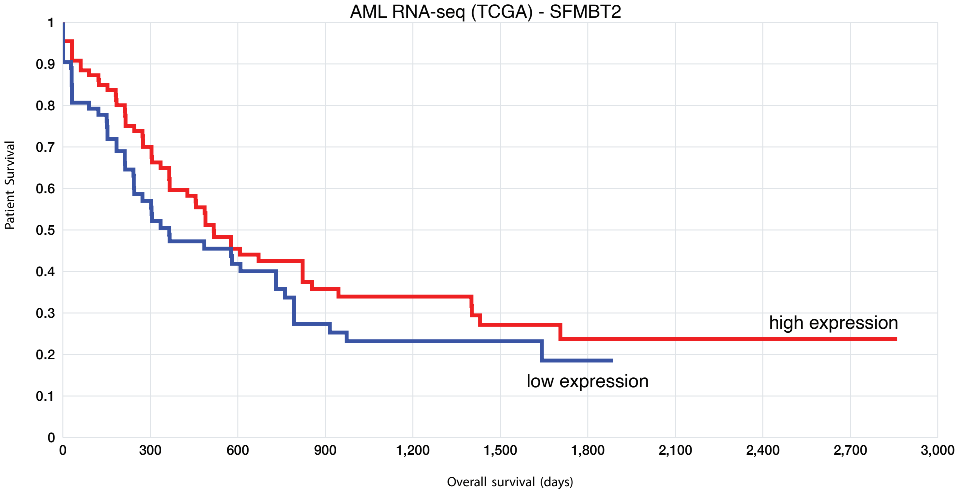

### Supplementary Figures (contd.)

#### S6. SFMBT2 in Normal Human Hematopoiesis

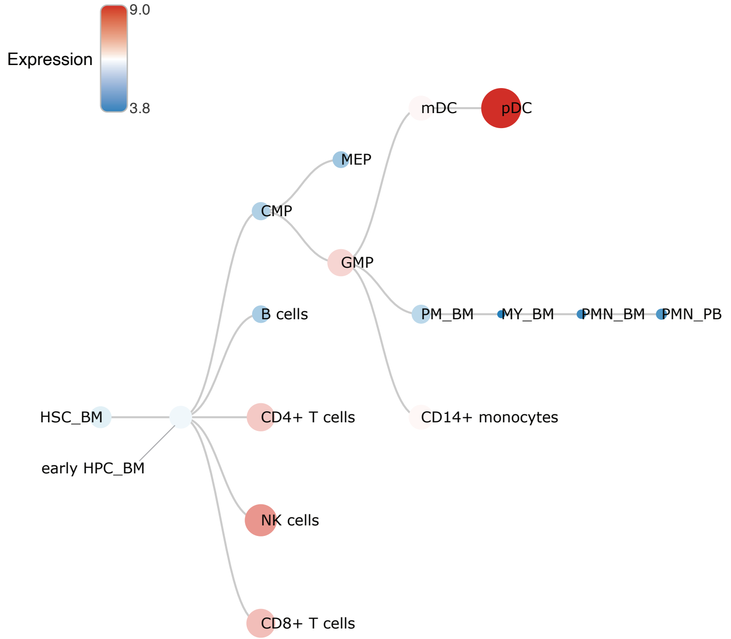


| Abbreviation | Name |
|---|---|
| HSC_BM | Hematopoietic stem cells from bone marrow |
| early HPC_BM | Hematopoietic progenitor cells from bone marrow |
| CMP | Common myeloid progenitor cell |
| GMP | Granulocyte monocyte progenitors |
| MEP | Megakaryocyte-erythroid progenitor cell |
| PM_BM | Promyelocyte from bone marrow |
| MY_BM | Myelocyte from bone marrow |
| PMN_BM | Polymorphonuclear cells from bone marrow |
| PMN_PB | Polymorphonuclear cells from peripheral blood |
| CD14+ monocytes | CD14+ Monocytes |
| B cells | CD19+ B cells |
| CD4+ T cells | CD4+ T cells |
| CD8+ T cells | CD8+ T cells |
| NK cells | CD56+ natural killer cells |
| mDC | CD11c+ myeloid dendritic cells |
| pDC | CD123+ plasmacytoid dendritic cells |

#### S7. SFMBT2 in Normal vs. AML Patient Hematopoiesis

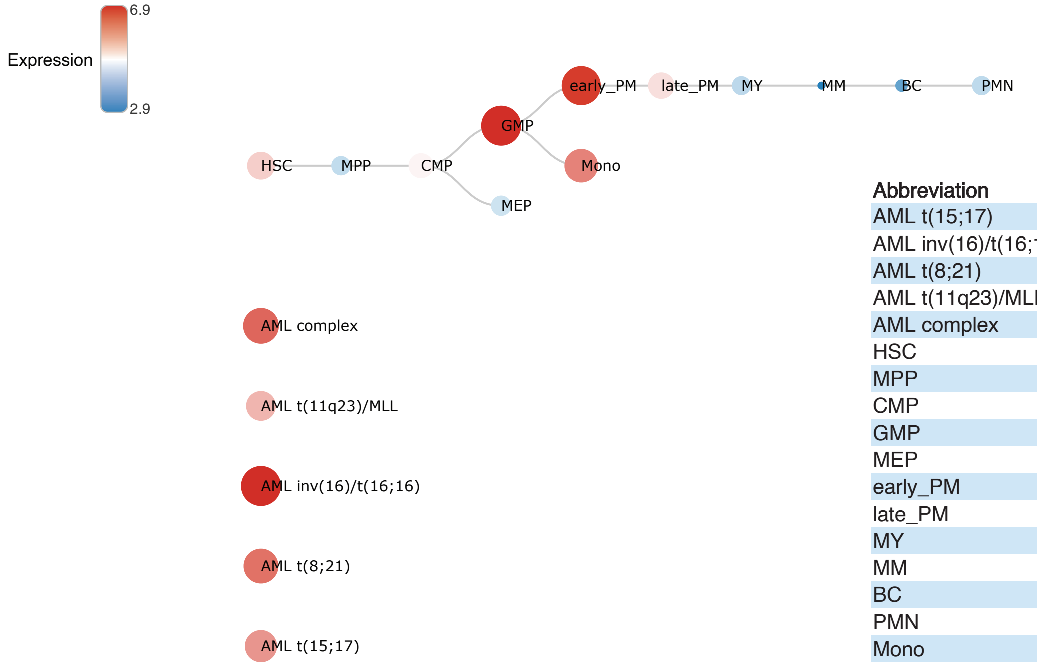


| Abbreviation | Name |
|---|---|
| AML t(15;17) | AML with t(15;17) |
| AML inv(16)/t(16;16) | AML with inv(16)/t(16;16) |
| AML t(8;21) | AML with t(8;21) |
| AML t(11q23)/MLL | AML with t(11q23)/MLL |
| AML complex | AML with complex aberrant karyotype |
| HSC | Hematopoietic stem cell |
| MPP | Multipotential progenitors |
| CMP | Common myeloid progenitor cell |
| GMP | Granulocyte monocyte progenitors |
| MEP | Megakaryocyte-erythroid progenitor cell |
| early_PM | Early Promyelocyte |
| late_PM | Late Promyelocyte |
| MY | Myelocyte |
| MM | Metamyelocytes |
| BC | Band cell |
| PMN | Polymorphonuclear cells |
| Mono | Monocytes |
